## Supplementary material for "Unbiased homeologous recombination during pneumococcal transformation allows for multiple chromosomal integration events": Reply to referees

Reviewer #1 (Minor Comments):

1. I only had one concern with the methodology- the role of the restriction modification (RM) systems is not resolved by these experiments. While the authors delete the restriction modification systems in the recipient cells and see no difference, the major issue with this is conclusion is that the donor tDNA is a PCR product, and hence not methylated, and hence not a target of the pneumococcal RM systems. I know pneumococcus can methylate donor DNA during transformation/uptake - but I would think for the authors to demonstrate no role for the RM systems, the donor DNA should have compatible/non-compatible methylation to fully support this conclusion.

I do not believe these experiments are needed to support the major conclusions- this could be rectified by changing the wording in the results section.

This is an excellent point and we have removed these experiments and this section.

2. Details with regards to the E. coli tDNA source used for these transformations- was this gDNA or PCR product? If PCR product, this section is fine as is but the experimental details need to be included in the methods.

It is PCR product. This has been clarified in the materials and methods section as below:
"For competition experiments using unrelated tDNA in **Figure 7**, DNA fragment that has no homology to pneumococcal genome but equal size (5 kb) and GC contents (~40% of GC) to *hlpA-mScarlet-I/spv_1159-sfGFP* fragments was amplified by PCR on genomic DNA of *E. coli* DH5alpha" (pages 37–38, lines 880–884)

3. Could the authors comment on the capture of extracellular DNA in the wild type background as opposed to the endA mutant? It would be helpful to serve a baseline and provide rationale for using this mutant in the measurement of surface bound DNA.

We have added a comment regarding the phenotype of wild type (*endA*$^{+}$) in the materials and methods section to clarify this as below:
"Analysis of DNA binding was performed in an *endA* mutant background (strain D39V *ssbB::luc (cm) endA::kan*), to favor accumulation of transforming DNA at the surface of competent cells. In wild type, *endA*$^{+}$ cells, surface-bound DNA is immediately internalized into the cytosol or degraded otherwise, which makes surface-bound DNA accumulation hard to be visualized as previously shown (page 36, lines 841–845).

4. I was unable to identify supplementary table S1 in the supplementary datasets- please ensure this is included with the final submission given the number of strains and genetic constructs utilized throughout this manuscript.

Table has been added

Reviewer #2 (General assessment and major comments (Required)):

In this work Kurushima et al. use recently developed fluorescent labelling techniques to study natural transformation in the human pathogen Streptococcus pneumoniae. Previously, genetic marker analyses have been used to study the different aspect of this process, but with these new techniques the process can now be studied at the single cell level. The authors used the single cell analysis to identify new transformation bottlenecks and tried to determine why some cells are genetically transformed and others are not. Related experiments have been performed in the past using classic genetics and Kurushima et al. were able to confirm these studies. In that sense, in my opinion, the novelty is limited and no important new molecular insights are provided. They found that the number of cells that are ultimately transformed is plateauing at approximately 50%, despite the fact that most cells bind DNA. This is partially the result of the heteroduplex formed after recombination followed by separation by strand replication, combined with the fact that the DNA binding sites on cells are limited so that there is a competition between DNA markers at saturating DNA concentrations. The authors argue that this mechanism entails a "fail-safe strategy for the population as half of the population generally keeps an intact copy of the original genome". I find this conclusion far-fetched for two reasons. Firstly, the DNA recombination event followed by DNA replication will automatically assure that only half the population will inherit the mutation, and to speak of a strategy implies that the organism has specifically evolved this system, but we are dealing here with a well-known and general recombination system found in many organisms that will generally result in a 50/50 distribution. Maybe more importantly, under natural conditions it is highly unlikely that cells encounter saturating levels of tDNA. In their experiments the authors use 3.2 nM DNA for transformation. If my calculation is correct, this would amount to 19xE11 DNA molecules per ml, which seems a bit high when assuming tDNA comes from lysed bacteria. In nature, this number will be much (much) smaller therefore there is no need for the bacterium to come up with a dedicated strategy to assure that not all cells in a population are being transformed. Finally, the results are very well presented and the paper makes easy reading.

We thank the reviewer for hinting to us to look at more environmentally relevant scenarios. To this end, we performed a set of new experiments (new Figure 8) in which we grew two pneumococcal strains together on agar plates in which one of the strains cannot become competent and carries SNPs conferring streptomycin and novobiocin resistance, respectively. In this scenario, horizontal gene transfer occurs via lysis of the recipient and by transformation of the then liberated environmental chromosomal DNA (new Figure 8A). One could argue this experiment more closely mimics the natural polymicrobial biofilm-like niche pneumococci encounter in the human nose, compared to our in vitro experiments using PCR products as donor DNA. As shown in new Figure 8B, also in this experimental setup, transformation is very efficient, up to $5.70 \times 10^{-2}$ (~0.05% of recipient cells are transformed) and markers transform independent of each other as demonstrated by our single cell analysis. We think this new experiment underscores that competence-based transformation is a main driver for pneumococcal genome evolution. We like to note that we do not think that the fail-safe has evolved as a dedicated strategy, but is rather a consequence of the recombination mechanism. This was mentioned in the Discussion as being a ‘spandrel effect’ and we have now further clarified this in the revised manuscript and changed fail-safe mechanism to fail-safe scenario.

Reviewer #2 (Minor Comments):

The only typo that I could find was on page 6 "work done by in the 1960".

It has been corrected as below:
“These findings correspond with work done in the 1960s and 1970s” (page 14, lines 326–327)

It is unclear what the control VL1800 in Fig. 6E should show.

Together with negative control (non-transformed VL1803), the positive control VL1800 strain shows that a proper threshold was used in our quantitative image analysis. Since even negative cells could still exert autofluorescence as a background and also even positive cells could attenuate the fluorescence signal, setting the proper threshold of fluorescence signal is crucial to quantify fluorescence signal acquired by transformation. Indeed, the fact that we found ~100% of double-positive or double-negative in VL1800 or non-transformed VL1803, respectively, showed that there is mostly no chance to misjudge fluorescence-positive cell as negative cell and vice versa.
To clarify this, we have added a sentence below in the figure legend of Fig. 6e and Fig. 7d:
"Analyzed data of the positive control strain VL1800 (*hlpA-mScarlet-I, spv_1159-sfGFP*) and negative control non-transformed strain VL1803 (*hlpA-stop-mScarlet-I, spv_1159-stop-sfGFP*) are also shown, demonstrating the accuracy of the threshold used to score positive transformants." (page 47, lines 1079–1082)
"Analyzed data of the positive control strain VL3126 (*ftsZ-mTurquoise2, spv_1159-msfYFP, hlpA-mScarlet-I*) and negative control non-transformed strain VL3127 (*ftsZ-stop-mTurquoise2, spv_1159-stop-msfYFP, hlpA-stop-mScarlet-I*) are also shown, demonstrating the accuracy of the threshold used to score positive transformants." (page 47, lines 1099–1102)

For the transformation test of the dpnI mutant (page 6), what DNA was used? I am not entirely sure, but I was of the impression that DpnI only cuts methylated DNA, so PCR DNA is not a good donor to test a dpnI mutant.

Good point, and also in reply to Reviewer 1, we have removed this section.

Reviewer #2 (Additional data files and statistical comments):

not applicable.

Reviewer #3 (Minor Comments):

1. Page 1, paragraph 1. Change "Streptococcus" to "Streptococci".

We have corrected the corresponding word to "Streptococci" as below:
"For example, one of the main genetic sources for penicillin resistance in *S. pneumoniae* is DNA acquired from non-pathogenic *Streptococci* from the viridans group such as *S. mitis* that also lives in the human nasal and oral cavities." (page 3, lines 45–48)

2. Page 2, right column. How do the amounts of the fluorescent-fusion proteins compare to the WT levels of these competence proteins and are the fusion proteins fully functional? Please comment briefly. Also, further down, some of the protein localizations look punctal and/or polar, rather than in bands. Is this expected, and if so, what are the implications of this localization pattern?

We like to note that we only used these fusions as reporters to estimate the proportion of cells that have activated the production of these machineries. Thus, for the current study it is not relevant whether they are functional or not and whether they are produced to wild type levels or not. Nevertheless, we tested their functionality (data not shown) and observed that ComGA fusion clearly impairs transformation efficiency compared to wild type. Both ComFA and ComEA fusions did not show significant effect on transformation efficiency compared to wild type. We believe that the punctual/polar localization patterns could be aggregated molecules since DNA is bound to the mid-cell region. Indeed, ComGA that shows strong punctual signal is unfunctional as mentioned above. Real localization of these proteins should likely be a band around mid-cell (Bergé, et al., 2013, PLoS Pathogens). Thus, we do not think the observations in this study is enough to discuss on functionality of those competence proteins at least based on sub-cellular localization and it is out of the scope for this study.

3. Page 4, right column, bottom. Change: "Upon take up" to "Upon uptake".

The corresponding sentence has been corrected as indicated.
"Upon uptake, integration and expression of exogenous transforming DNA (tDNA) containing ~" (page 9, lines 186–187)

4. Page 5, bottom paragraph. The heading of this section could be changed to make it more descriptive of the main conclusion.

Good point. The heading has been changed to "Single cell quantification of homeologous recombination highlights transformation bottlenecks" (page 10, lines 230–231)

5. Page 6, bottom, left column. Change "Fig. S7" to "Fig. S8".

"Fig. S7" has been corrected to "Fig. S8 (**Figure 4–figure supplement 2, now**)" in corresponding line. (page 12, line 280)

In top of next column, cite Fig. 4d.

Fig. 4d has been cited as below:
"To test if our transformation reporter system depends on the competence-induced homologous recombinase, RecA, we depleted RecA expression level using CRISPR interference (Figure 4D)." (page 12–13, line 280–282)

6. Page 6, last paragraph. Increase the size of Fig. S11b. It's tiny and hard to read.

Size of Fig. S11b (Figure 5–figure supplement 2, now) has been increased in size.

Cite Fig. 5 in the sentence that begins with "balk".

Fig. 5 has now been cited as below:
"To test whether the genomic location of the recombination site and the read orientation of the newly acquired functional allele influences transformation efficiency, we introduced the *spv_1159-stop-sfGFP* reporter at 4 different positions on the chromosome: on the right arm of the chromosome at 101° of the circular chromosome (*bgaA* locus), near the terminus at 169° (*hlpA* locus), on the left arm of the chromosome at 295° (*cep* locus) and near *oriC* at 359° (*comCDE* locus) (Figs. 5A and B). In addition, *spv_1159-stop-sfGFP* was introduced on both

the positive and negative strand on the left arm of the chromosome (*cep* locus at 295°) and on the right arm of the chromosome (*bgaA* locus at 101°) (Fig. 5C).” (page 13, lines 296–299)

Correct typo "work done by in".

It has been corrected as below:
“These findings correspond with work done in the 1960s and 1970s” (page 14, lines 326–327)

At end, "dsDNA transformation events" is shorthand, but perhaps misleading. Please consider changing.

Original sentence "dsDNA transformation events" has been changed to “double transformation events on both strands”. (page 15, line 334)

7. Page 7, first sentence. "Data" is plural. Please correct here and check throughout.

They have been corrected.

8. In Figure 5 and Figure 6 and other places. There is really no sense of how many independent biological replicate experiments are included in the data sets.

Experiments pointed out here are represented from one experiment except for histograms showing multiple transformation outcome, but the experiments have been done at least 3 times with similar results and representative outcomes are shown. This is now clarified in the figure legends.

In addition, in the right panel, the interpretation is that there is no effect of strand orientation on transformation efficiency; yet, the frequency numbers for the experiment shown vary between 22% to 44%. Is this difference not significant in this experiment? Please clarify this point.

This is a good point. We do think that the difference between 22% and 44% is significant, but that this is due to the chromosomal position effect as briefly discussed. However, there is no significant difference in the + or – strand at the same locus. This has now been clarified in the revised manuscript.

Likewise, in Figure 6e, does this histogram represent the results of one experiment or multiple replicate experiments. Please clarify.

Number of replicates of histograms in Fig. 6e (and Fig. 7d as well) is described in the figure legend as below:
“Bars represent mean ± SD of three independent replicates.” (page 46–47, lines 1078–1079 and page 47, line 1098–1099)
